# Taxonize-gb: A tool for filtering GenBank non-redundant databases based on taxonomy

**DOI:** 10.1101/2024.03.22.586347

**Authors:** Mohamed S. Sarhan, Michele Filosi, Frank Maixner, Christian Fuchsberger

**Affiliations:** Institute for Biomedicine, Eurac Research, Bolzano 39100, Italy (Affiliated institute with Lübeck University, Lübeck, Germany); Department CIBIO, University of Trento, Trento 38123, Italy; Institute for Mummy Studies, Eurac Research, Bolzano 39100, Italy

## Abstract

Analyzing taxonomic diversity and identification in diverse ecological samples has become a crucial routine in various research and industrial fields. While DNA barcoding marker-gene approaches were once prevalent, the decreasing costs of next-generation sequencing have made metagenomic shotgun sequencing more popular and feasible. In contrast to DNA-barcoding, metagenomic shotgun sequencing offers possibilities for in-depth characterization of structural and functional diversity. However, analysis of such data is still considered a hurdle due to absence of taxa-specific databases. Here we present taxonize-gb, a command-line software tool to extract GenBank non-redundant nucleotide and protein databases, related to one or more input taxonomy identifier. Our tool allows the creation of taxa-specific reference databases tailored to specific research questions, which reduces search times and therefore represents a practical solution for researchers analyzing large metagenomic data on regular basis. Taxonize-gb is an open-source command-line Python-based tool freely available for installation at https://pypi.org/project/taxonize-gb/ and on GitHub https://github.com/msabrysarhan/taxonize_genbank. It is released under Creative Commons Attribution-NonCommercial 4.0 International License (CC BY-NC 4.0).

## Introduction and motivation

Environmental metabarcoding is a powerful molecular biology technique used to analyze the biodiversity within complex environmental samples (1). It operates by targeting and amplifying specific DNA regions, such as the 16S ribosomal RNA gene for bacteria, the COI gene for animals, or trnL for plants from a mixed sample of organisms. Once amplified, these genetic sequences are then subjected to high-throughput DNA sequencing, generating millions of short DNA sequences. Custom bioinformatic tools should be developed to in silico match these sequences to reference databases, allowing to identify and quantify the different species present in the original sample (2). Environmental metabarcoding has been used in different areas of research. For example, it has been used in food authentication (3), to understand plant-pollinator interactions (4), to detect invasive species in the environment (5), to monitor anthropogenic pollution (6), to reconstruct dietary components (7), and to reconstruct ancient ecosystems (8).

Most of these studies relied on targeted amplicon approach, which employs amplification of single-marker genes to target specific taxa (9,10). However, choosing the appropriate marker gene for each taxon remains a perplexing issue, given the varying sensitivity and resolution levels of different markers. Selection of maker genes is also highly dependent on the quality of the reference database and availability of suitable unbiased universal primers, which ends up in a trade-off situation between feasible in vitro amplification and reliable in silico identification. Therefore, various studies suggested usage of multiplexed marker genes, which reported to be efficient in increasing species detection (11,12). However, such approach doubles the overall computational cost of the analysis which adds another factor to be considered in the trade-off.

During the past decade, the costs of next-generation sequencing (NGS) have continued to decrease, making it more affordable for all biology disciplines. Such affordability encouraged environmental DNA researchers to shift towards the use of shotgun metagenomic sequencing instead of targeting only single or few marker gene amplicons. Shotgun metagenomic sequencing offers multiple advantages to the environmental DNA, such as targeting more genomic regions and avoiding the PCR bias-related issues. Using shotgun metagenomics is sometimes an indispensable approach particularly when analyzing very low biomass and low DNA samples, like in the cases of ancient DNA analysis or forensics, where the DNA is highly damaged which makes the retrieval of DNA amplicons a highly challenging task and amenable to many technical biases (13).

While several curated databases have been already established to provide valuable resources for metabarcoding marker genes, a recurring challenge has been the lack of consistent maintenance and updates. These databases often start off as well-structured repositories of accurate and organized data, which over time with the rapid pace of new deposited data, the information contained within these databases become outdated. There are some well curated marker gene databases that are regularly updated and maintained (14), however they do not include all used metabarcoding marker genes. Therefore, there are new initiatives to develop database curation tools, such as Bcdatabaser (15) MetaCurator (16), which allow the user to develop up-to-date custom databases suitable for specific research questions and confined to particular taxonomic groups of interest.

While the analysis of shotgun metagenomic data requires more comprehensive genome-wide databases, it also requires high computational resources and often specialized infrastructures, to handle such big data. Therefore, selection of targeted curated non-redundant and up-to-date databases is of paramount importance. Accordingly, the National Center for Biotechnology Information (NCBI) hosts GenBank, which contains a comprehensive collection of genetic sequence data, including DNA, RNA, and protein sequences submitted by researchers from around the globe (17). The NCBI offers up-to-date non-redundant protein and nucleotide databases (18), which seem to be the most suitable reference databases for analyzing shotgun sequences from environmental DNA samples (19). However, due to their comprehensiveness and regular updates, there are two major concerns. First, these databases grow exponentially every year which makes them difficult to maintain even with big computational infrastructures (18). Second, they contain a lot of off-target references which are impractical to keep in the search database, especially when the researcher is interested in specific taxonomic group. For example, if the researcher is interested in analyzing plant diversity, it would be a waste of resources to keep all non-plant proteins/nucleotides in the search database (e.g., animals, bacteria, phages, etc.).

Using taxa specific databases as reference to analyze shotgun metagenomic sequences could help in detangling this issue. However, such specific databases are not offered by the GenBank nor by other genomic repositories. Although the GenBank is offering now an online experimental BLAST non-redundant nucleotide database on domain level (Eukaryotes, Prokaryotes, and Viruses), the sequences of these databases are not available for download for offline command line usage.

This issue becomes more pronounced when dealing with eukaryotic diversity, since in contrast to bacterial and archaeal diversity analysis, there are not many tools which are optimized for their analysis.

### Software functionality

We developed the software tool “taxonize-gb” as a command-line tool, developed in Python 3, designed to streamline the retrieval and filtering of data from the NCBI GenBank protein and nucleotide databases. The tool comprises various modules, with one specifically tailored for accessing the File Transfer Protocol (FTP) directories of the NCBI (https://ftp.ncbi.nlm.nih.gov/) to retrieve the GenBank database files, i.e., nt/nr FASTA-formatted sequence files, mapping of accession numbers to taxonomy IDs, and the NCBI taxonomy database (20).

The core module of our tool is “taxonize_gb” which is designed to streamline data extraction from the NCBI GenBank NR/NT databases based on a user-specified taxonomy IDs (TaxID). The module “taxonize-gb” performs filtering on the non-redundant protein/nucleotide databases of the GenBank based on a specified TaxID, which can be at any taxonomic level. The module performs the filtering on three main steps: (1) It employs the module DiGraph of NetworkX (21) to parse content of the “nodes.dmp” and the “names.dmp” files from the taxonomy database, representing them as a graph structure. Then, based on the user provided TaxID, it extracts all descendant TaxIDs (graph nodes) and outputs them as a data frame to store them along with their corresponding scientific names (**Figure 1**). (2) The module filters the mapping files (i.e., accession to taxonomy ID) to retain only the accession numbers associated with the input TaxID and its descendants. (3) In the last step, it uses the Biopython modules (22) to parse the non-redundant FASTA sequences and perform a search within their headers to identify the filtered accession numbers, and optionally user-provided keywords (**Figure 1**). The module is designed to take minimal and flexible inputs from the user – The user can provide paths for different database files; in case they are available in the local system. For example, to get all non-redundant Viridiplantae protein records, you can run the following command:

**Figure 1.**
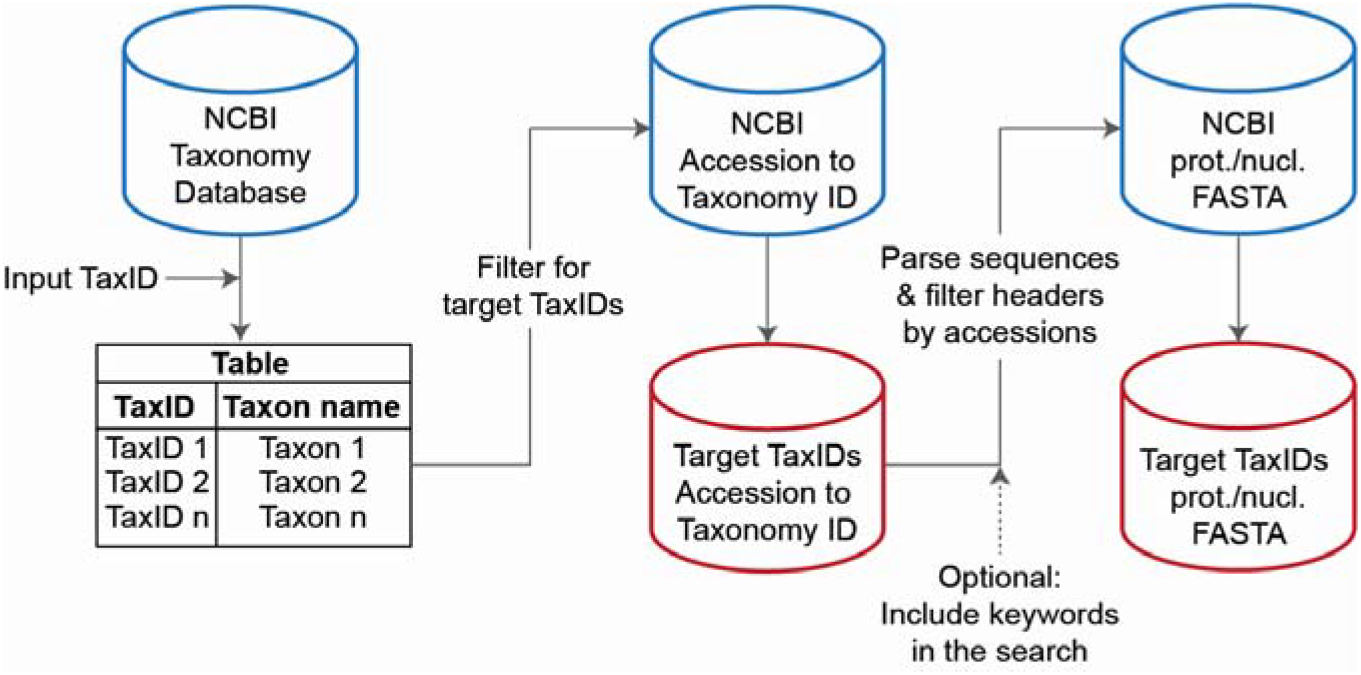
Visual workflow for the “taxonize_gb” module for filtering the NCBI non-redundant protein and nucleotide databases.

~~~
taxonize_gb --db nr –db_path nr.gz --prot_acc2taxid
prot.accession2taxid.gz --pdb_acc2taxid pdb.accession2taxid.gz --
taxid 33090 --out Viridiplantae_nr/
~~~

While if the user does not provide any of the input databases, the latest version of the necessary databases will be downloaded automatically based on the provided mandatory input ‘--db’ option. For example, to get all non-redundant Viridiplantae protein records, the user can run the following command:

~~~
taxonize_gb --db nr --taxid 33090 --out Viridiplantae_nr/
~~~

Additionally, the module can utilize the optional features of including keywords in the search to refine the filtering, e.g., to filter for specific gene/protein names or to filter for organellular genes/genomes. Detailed explanations on how to use the modules with further examples are available in the GitHub repository page (https://github.com/msabrysarhan/taxonize_genbank).

The last module in our tool is “get_taxonomy” which is a utility script that uses the ete3 toolkit to retrieve taxonomic lineages of a given FASTA file. This module would be useful when the user is interested to make an overview on the taxonomic distribution of the filtered databases.

### Evaluation

The alternative available option to perform taxa-specific search using the NCBI non-redundant protein database (to the best of our knowledge) is to use DIAMOND search tool against the complete NCBI-nr database, restricting the search to specific taxonomic IDs (using “--taxonlist” flag).

To evaluate the performance, we explored the efficiency of different search approaches for querying NCBI non-redundant protein database (NCBI-nr). We compared two methods of DIAMOND search: against taxonized NCBI-nr databases and against complete NCBI-nr database with restricted TaxIDs. For the comparison, we targeted the following taxa and TaxIDs: 1) Chordata [taxid: 7711]; 2) Fungi [taxid: 4751]; and Viridiplantae [taxid: 33090]. We used the published metagenomic data from ancient paleofeces (23). For each search job, we used 16 CPUs and 50 Gb of RAM.

Our results clearly demonstrate a substantial advantage in terms of time efficiency when employing DIAMOND search against taxonized GenBank databases (**Figure 2**). The search times were significantly shorter when using taxonized databases (means 1.13 - 1.45 h), as opposed to the traditional complete database searches with TaxIDs restrictions (means 8.96 - 10.8 h).

**Figure 2:**
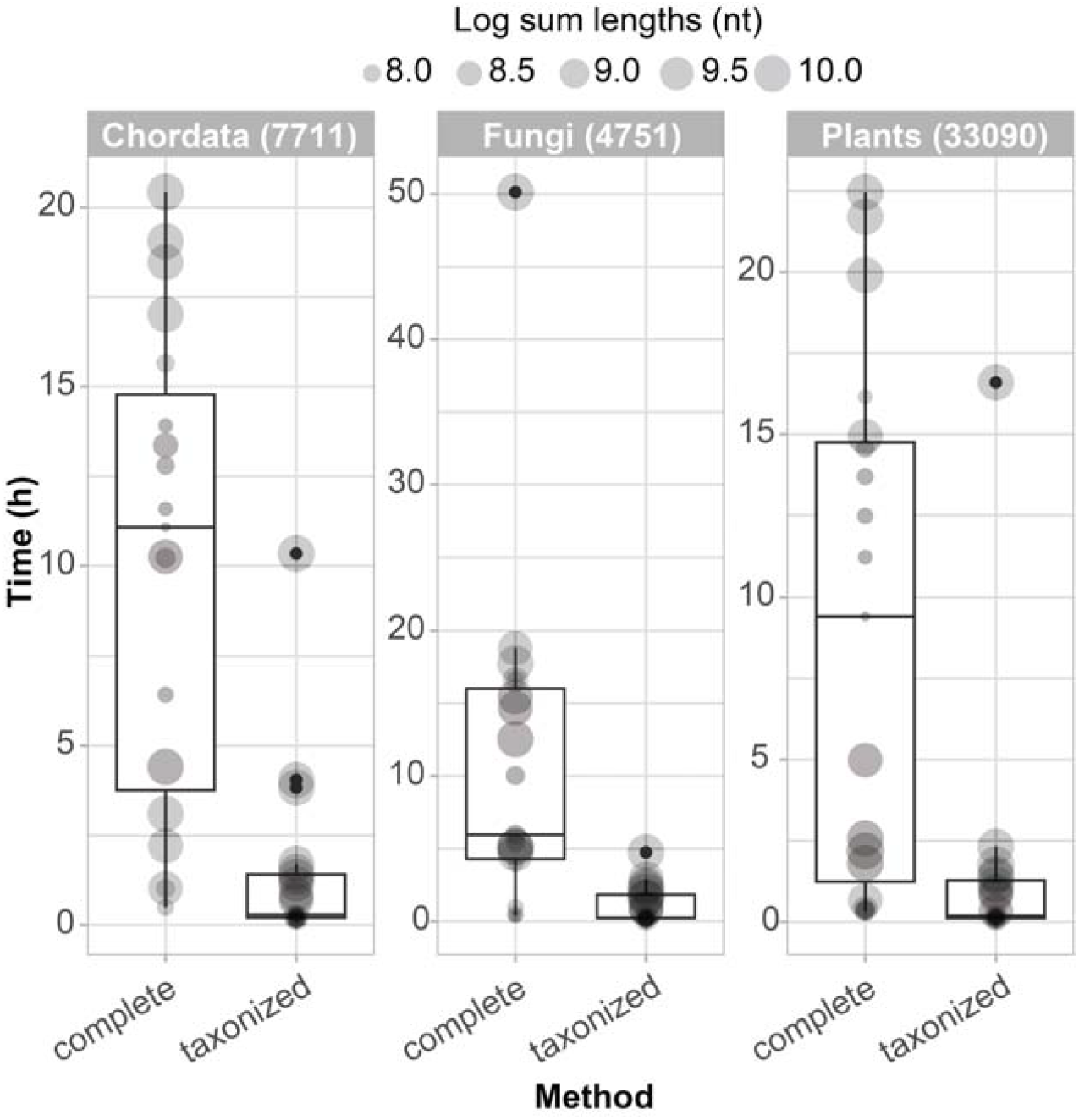
Performance comparison in terms of the runtimes (h) of DIAMOND search against taxonized NCBI-nr vs complete NCBI-nr with restricted TaxID search option. For further information on the used metagenomic samples, please refer to Maixner et al. (2021). The data are publicly available at ENA: PRJEB44507.

## Conclusion

Taxonize-gb is a versatile, easy-to-use command-line tool that provides a comprehensive solution for researchers working with metagenomic data across multiple research disciplines by enabling efficient downloads and taxonomy-based filtering. This gives researchers the flexibility to focus their analyses on their area of interest. Finally, the reduced search times provided by using taxonized databases provide a practical and beneficial solution for researchers dealing with large metagenomic data and data-intensive research projects.

## Acknowledgements

We are grateful to the support of the Life Science Compute Cluster (LiSC) of the University of Vienna. We thank Mohamed R. Abdelfadeel of Leibniz-IGZ for testing the tool.

